# Spatio-temporal analysis of the innate immune response to cytoplasmic dsDNA using a novel cGAMP biosensor

**DOI:** 10.1101/2024.06.10.598238

**Authors:** S Smarduch, SD Moreno-Velasquez, S Dadsena, D Ilic, R Morant, G Pereira, M Binder, AJ García-Sáez, SP Acebrón

## Abstract

The cGAS-STING signalling pathway acts as gatekeeper in the innate immune response towards extrinsic and intrinsic sources of cytoplasmic double stranded DNA (dsDNA). At the core of this pathway is the cGAS-dependent production of the intra- and extra-cellular messenger 2’3’-cyclic GMP-AMP (cGAMP), which activates STING and leads to IRF3-dependent expression of cytokines and interferon. Despite its relevance to monitor viral and bacterial infections, cell death, and genome instability, the lack of specific live reporters has precluded spatio-temporal analyses of cGAS-STING signalling. Here, we generate a fluorescent cGAMP biosensor by engineering the functional interaction of activated STING and IRF3 at the Golgi. We show that cells encoding for the cGAMP biosensor react within 45 minutes, and in a time- and concentration-dependent manner to STING agonists and cGAMP. We demonstrate that the cGAMP biosensor is suitable for single cell characterisation of the dynamics of Herpes Simplex Virus 1 infection, mtDNA release upon apoptosis, and other sources of cytoplasmic dsDNA. Intriguingly, we show that STING signalling is not activated by ruptured micronuclei, suggesting that other cytosolic pattern recognition receptors underlie the interferon response upon chromosomal instability. Furthermore, we demonstrate that the biosensor sensitivity is sufficient to report microenvironmental cGAMP, allowing to analyse how STING signalling spreads through neighbouring cells, including upon viral infection. Taken together, we provide a tool to easily monitor the activation cGAS-STING signalling pathway by live cell imaging, plate reader or immunofluorescence analyses; and to capture the spatio-temporal and heterogenous dynamics of the response to cGAMP.

## INTRODUCTION

The cGAS-STING signalling pathway has been proposed as a versatile and unifying sensing mechanism for double stranded DNA (dsDNA) that is coupled to inflammation in mammals ^1-3^. The catalytic activity of cGAS is stimulated by double stranded DNA (dsDNA) ^4-6^ from any source, and therefore uncoupled from pathogen- or host-specific features ^3^. Mechanistically, a positive charged nucleotidyl-transferase domain binds the dsDNA backbone thereby inducing allosteric changes that favour ATP and GTP binding, and the subsequent synthesis of 2’3’-cyclic GMP–AMP (cGAMP) ^6^. Dimerization of cGAS results in a 2:2 DNA-cGAS complex and a ladder-like structure along the dsDNA backbones ^7-9^. In vitro analyses demonstrated that longer dsDNA fragments (over 4 kbp) are more efficient activators of cGAS, possibly due to emerging properties associated to phase-separation ^9,10^.

STING is an endoplasmic reticulum (ER) resident transmembrane protein. STING consists of a lumen N-terminal domain, 4-span transmembrane domains, and a cGAMP ligand binding domain (LBD) facing the cytoplasm and bound to a C-terminal tail ^11^. In the absence of cGAMP, STING forms homodimers with the ligand binding domains interlinked and covered by their respective C-terminal tails. STING-cGAMP complex formation induces allosteric changes that i) lead to additional oligomerisation between activated dimers via bisulphite bridges and ii) presentation of the C-terminal tails ^2,11-14^. Activated STING is trafficked to the Golgi via the COPII machinery in a process that is not yet well-understood ^15,16^. At the Golgi, STING is palmitoylated, which is essential for its activity ^15^. Upon activation, the C-terminal tails of the STING complexes recruit the TANK Binding Kinase 1 (TBK1), which dimerises through auto-phosphorylation. TBK1 also phosphorylates the C-terminal tails of STING ^17^, which is required for the subsequent recruitment and phosphorylation of the interferon regulatory factor 3 (IRF3) ^18^. Phosphorylated IRF3 dimerises and translocates to the nucleus where it induces the transcription of type I interferon genes, interferon-stimulated genes, and inflammatory cytokines ^19^.

In the context of host DNA, both mitochondrial DNA (mtDNA) during apoptosis, as well as genomic DNA (gDNA) resulted from DNA damage or chromosome missegregation can induce cGAS-STING signalling. During apoptosis, the pro-death factors BAK and BAX oligomerise at the mitochondrial outer membrane ^20^, leading to its permeabilization and the release of cytochrome C and mtDNA to the cytoplasm ^20,21^. Interestingly, cytochrome C signalling leads to caspase activation that can attenuate cGAS-STING signalling by cleaving of cGAS, TBK1 and IRF3 to ensure that the immunogenic response is not activated during apoptosis ^22^. In the absence of caspase-signalling, the efflux of mtDNA fully activates cGAS-STING signalling ^20,22,23^. Missegregated chromosomes, broken gDNA fragments arisen from chromosome bridges, or otherwise damaged DNA, are normally encapsulated by their nuclear envelope in the daughter cells in so-called micronuclei ^24,25^. Damaged DNA at the micronuclei undergoes chromothripsis (extreme rearrangements and pulverisation) due to the collapse of the lamina-defective nuclear envelope ^25^. This process exposes the DNA to the cytoplasm leading to the accumulation of GMP–AMP synthase (cGAS) in the micronuclei and the subsequent activation of STING signalling ^24,26^. Of note, an inhibitory phosphorylation by CDK1 and AurB ^27,28^, as well as membrane tethering ^28^, prevents cGAS activation by condensed chromosomes during a normal mitosis, although it can allow slow accumulation of IRF3 and subsequent apoptosis upon mitotic arrest ^29^.

In the context of foreign DNA, viruses, bacteria, as well as gDNA/mtDNA released from death cells can trigger the activation of cGAS-STING. For instance, infection of macrophages by Herpes Simplex Virus 1 (HSV-1) leads to a quick response towards the viral DNA by cGAS ^30^, which triggers interferon response necessary for the clearance of the infected cells. Similarly, intracellular replication of the gram-positive bacterium *Listeria monocytogenes* triggers cGAS-STING activation and interferon expression in myeloid cells ^31^.

Intracellularly accumulated cGAMP can be exported by ABCC1 transporters to the microenvironment ^32^, where it functions as immune-transmitter ^33^ imported by SLC19A1 and SLC46A2 ^34,35^, including by macrophages and monocytes. However, this immune response can be hampered by the ecto-nucleotide pyrophosphatase/phosphodiesterase ENPP1 ^36^, which degrades extracellular cGAMP ^37^.

Although several transcriptional reporters to cascades downstream of STING have been used in the past to, including the INFb-Luciferase reporter, these approaches have a long latency on their response – often conflicting with cell death –, lack spatio-temporal information, and can be targeted by other signalling cascades, independently of STING. As such, the study of the innate immune response in live cells is hampered by the lack of robust biosensors and reporters. The STING receptor in particular, acts as a bottleneck in the surveillance of any source of cytoplasmic dsDNA. As such, engineering STING as biosensor could provide an opportunity to monitor many different inputs ranging from chromosomal stability, viral/bacterial infections, apoptosis and other forms of cell death, auto-immunity and tumour immunity, including in cancer therapy.

## RESULTS

### Design and validation of a novel cGAMP biosensor

We turned our attention towards the mechanisms of STING activation and designed different chimeric proteins using cGAS, STING, TBK1, IRF3, NFκβ, and IKK, as well as linkers of various lengths, to couple their activation or interaction with luminescence, CRISPRa, and fluorescence reporters. Among the tested candidate constructs, we selected a tandem split GFP approach between STING and its downstream target IRF3 (Figure 1A,B). In detail, we generated a construct containing STING-GFP11_x3_ (GFP(N)) and IRF3-GFP1-10 (GFP(C)) separated by a ribosome skipping peptide (P2A) to facilitate stochiometric expression (Referred from here on as cGAMP biosensor). To avoid the constitutive activation of STING signalling due to plasmid transfection of the cGAMP biosensor and to study its function in different cell models, we generated HEK293T, HeLa, and human fibroblasts stable cell lines using the PiggyBac transposase system. Western blot analyses of cGAMP biosensor HeLa cells upon treatment with the STING agonist diABZI induced phosphorylation of STING-GFP(N) and the endogenous downstream effector TBK1 (Figure 1C), confirming the functionality of the modified receptor.

**Figure 1:**
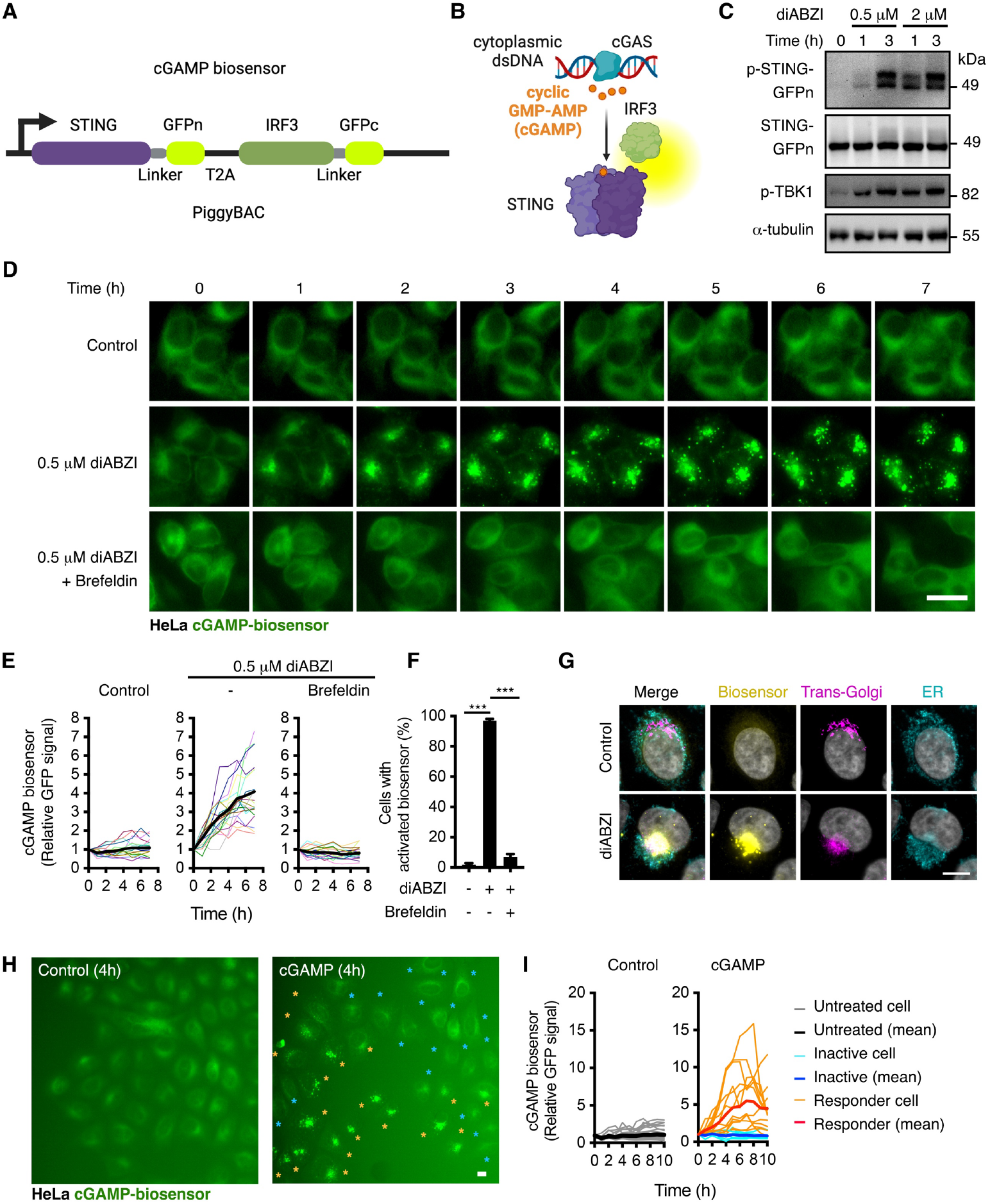
Design and validation of the cGAMP biosensor. **A**,**B**, Schematics of the PiggyBAC construct containing the cGAMP biosensor (**A**) and the suggested mode of function (**B**). **C**, Western blot analyses in HeLa cells stably transfected with the cGAMP biosensor upon treatment with the STING agonist diABZI. **D-F**, Live cell imaging analyses of HeLa cGAMP biosensor cells in the presence or absence of diABZI and the ER to Golgi transport inhibitor Brefeldin A. Note that each coloured line is one single cell randomly selected from the population. In (**F**) activation of the biosensor was scored as clustering of the GFP signal in 1 or more puncta, and it is represented as mean ± SD of > 200 cells per condition. **G**, immunofluorescence co-localization analyses of HeLa cGAMP biosensor cells in the presence or absence of diABZI for 4 hours. **H**,**I**, Live cell imaging of HeLa cGAMP biosensor cells in the presence or absence of 42 μM cGAMP. In (**I**), cells responding within the first 4 hours (49%) are shown in orange while non-responding cells (51%) are shown in blue. *P*-values from one-way or two-way ANOVA between independent experiments from the indicated groups are indicated as *P < 0.05, **P < 0.01, ***P < 0.001, or n.s., not significant.

Live cell imaging of the cGAMP biosensor in HeLa cells showed a diffuse EGFP signal in the ER (Figure 1D). Treatment with the STING agonist diABZI (0.5 μM) led to quick clustering of the cGAMP biosensor within ∼1 hour (Figure 1D-F), reaching a 4-fold increase in fluorescence at 8 hours. Consistent with its requirement for STING-IRF3 interaction, inhibition of ER to Golgi transport via co-treatment with Brefeldin A, fully blocked the activation of the biosensor in HeLa cells (Figure 1D,E). Of note, virtually all cells harbouring the biosensor (97%) responded to 0.5 μM diABZI within the first 4 hours, while only 2% of the controls displayed activation of the biosensor (Figure 1F). Immunofluorescence analyses confirmed the co-localisation of the inactive cGAMP biosensor at the ER (Figure 1G). The activated biosensor localised in large vesicles around the Trans-Golgi, consistent with a reported role of endolysosomal trafficking in STING signalling termination ^38^. Live cell imaging analyses of HEK293T cells and fibroblasts (HFF-1) stably expressing the cGAMP biosensor and treated with diABZI were largely consistent with the studies in HeLa cells (Figure S1A,B).

Next, we examined the response of the biosensor directly to microenvironmental cGAMP, without cell permeabilization, thereby relying on its import by solute carriers ^34,35^. Unlike the cell permeable STING agonist diABZI, incubation with 42 μM cGAMP led to a heterogenous response of the biosensor in HeLa cells, with ∼50% of cells displaying activation of the receptor at different amplitudes and dynamics within the first 4 hours (Figure 1H,I), while the other half remained inactive. These results suggest that cell intrinsic (e.g., cell cycle, stochastic expression of signalling modulators) or extrinsic (e.g., confluency, expression of inhibitors) can limit the import and/or cellular accumulation cGAMP.

Titration experiments in HeLa cells with diABZI showed a similar time- and concentration-dependent response of the cGAMP biosensor comparing with phospho-STING and phospho-TBK1 analyses by Western blots, but instead providing single cell resolution (Figure 2A-F). In particular, live cell imaging analyses showed that the biosensor clustered in HeLa cells treated with as little as 50 nM diABZI (Figure 2A,B) and ii) 78% of HeLa cells showed activation of the biosensor after just 45 min exposure to 2 μM diABZI (Figure 2D,E). To examine the recovery of the biosensor HeLa cells were first incubated for 6 hours with 0.5 μM diABZI and imaged upon withdrawal of the STING agonist. The activated biosensor was largely cleared within 20 hours from the cells, without impairing growth and further expansion of the cells (Figure 2G and not shown). To compare the biosensor with the downstream response of STING signalling we turned into HEK293T cells, which unlike HeLa cells, lack cGAS/STING but display a functional target gene response. As expected, diABZI only induced reliable luciferase activity after 24 hours in IFNb-luc reporter assays (not shown). More importantly, HEK293T expressing the cGAMP biosensor were able to transduce signalling and lead to 6-fold activation of the reporter upon 2 μM diABZI, similarly to HEK293T cells expressing wt STING (Figure S1C). To determine the suitability of the cGAMP biosensor for large scale analyses, we performed siRNA assays in 384-well format using siRNA against TBK1 as positive control and imaging the plates in a large content microscope. Consistent with its requirement for STING activation, knock down of TBK1 blocked the activation of the biosensor upon treatment with diABZI (Figure 2H-J).

**Figure 2:**
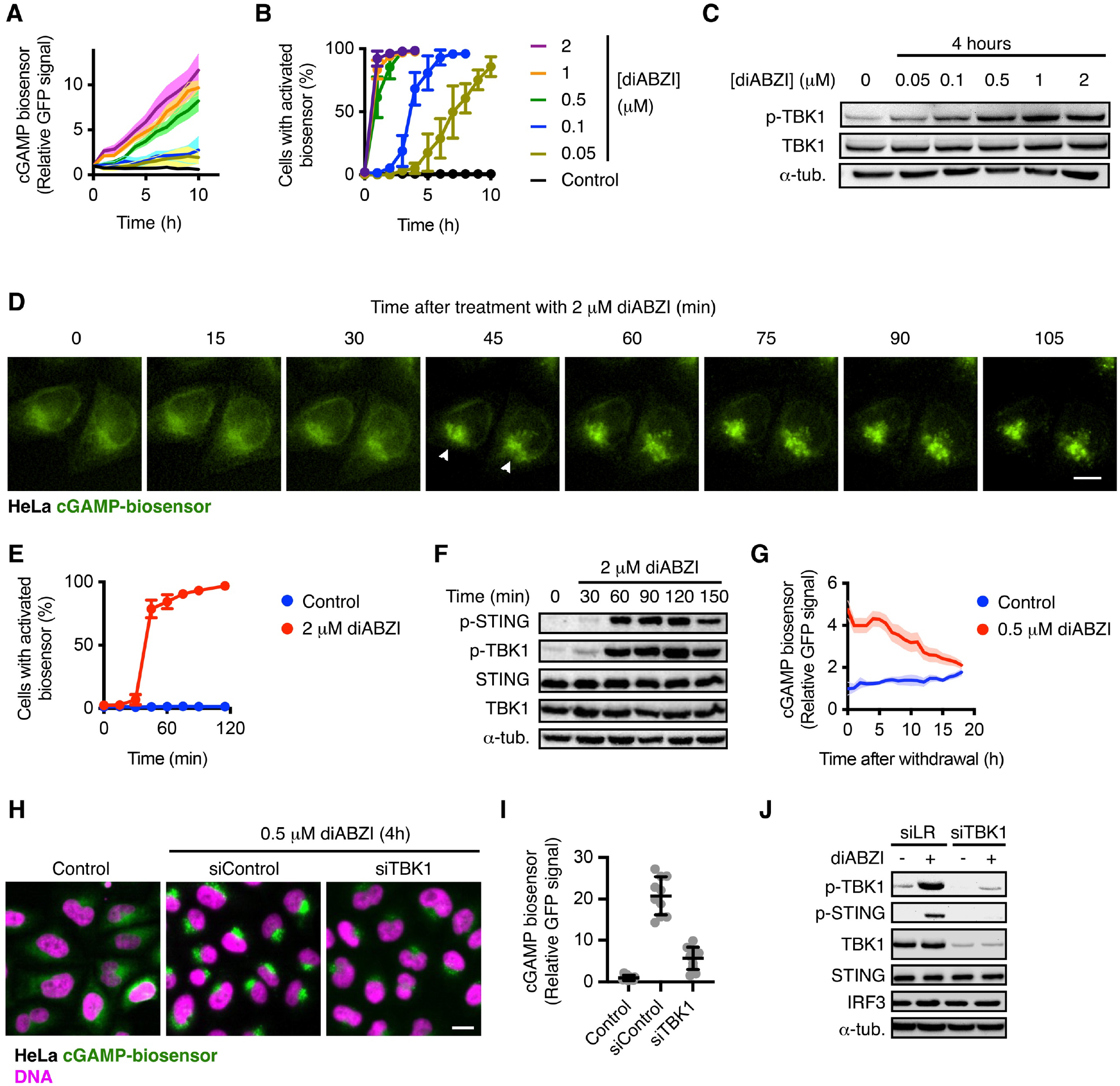
Characterisation of the cGAMP biosensor dynamics. **A**,**B**, Live cell imaging analyses of HeLa cGAMP biosensor cells in the presence of the indicated concetrations of diABZI. In (**A**) each condition is showed as the mean ± SED of 20 randomly selected cells. In (B) activation of the biosensor was scored as clustering of the GFP signal in 1 or more puncta, and is showed as mean ± SD of > 200 cells per condition. **C**, Western blot analyses in wt HeLa cells upon treatment with the STING agonist diABZI. **D**,**E**, Live cell imaging of HeLa cGAMP biosensor cells in the presence or absence of diABZI. In (**E**) activation of the biosensor was scored as clustering of the GFP signal, and it is represented as mean ± SD of > 200 cells per condition. **F**, Western blot analyses in wt HeLa cells upon treatment with the STING agonist diABZI. **G**, Live cell imaging of HeLa cGAMP biosensor cells upon withdrawal of diABZI. **H**,**I**. immunofluorescence analyses of HeLa cGAMP biosensor cells transfected with the indicated siRNAs and treated as indicated for 4 hours. Note that cells were imaged and analysed in a 384x high content microscope prepared for screening analyses. **J**, Western blot analyses in wt HeLa cells upon transfection with unrelated control siRNA (siLRP6; siLR) or siTBK1 and treatment with the STING agonist diABZI. *P*-values from one-way or two-way ANOVA between independent experiments from the indicated groups are indicated as *P < 0.05, **P < 0.01, ***P < 0.001, or n.s., not significant.

Taken together, these results show that the cGAMP biosensor monitors the STING signalling response, including receptor activation and transport from ER, as well as TBK1 and IRF3 recruitment, and provides a fast (45 min), reversible, and single cell resolution readout to STING agonists and microenvironmental cGAMP.

### Monitoring foreign dsDNA with a novel cGAMP biosensor

Next, we examine the response of the biosensor to foreign sources of dsDNA. Transfection of plasmid DNA expressing mCherry induced the activation of the cGAMP biosensor in 95% of the HeLa cells that would become mCherry positive (transfected) during the following 24 hours of live-cell imaging (Figure 3A,B). Of note, plasmid DNA transfected cells displayed a largely synchronised response starting 4 hours post-transfection, which peaked at 8 hours (Figure 3A,C). Interestingly, 44% of the untransfected neighbouring cells (mCherry negative during at least 24 hours) activated their biosensors, albeit with ∼2-hour median delay compared to plasmid DNA carrying cells. Further analyses did not reveal a mCherry^+^ distance-dependent activation in the untransfected cells (mCherry^-^), suggesting –as in the case of the direct cGAMP treatments (Figure 1H,I)– other factors regulate the stochastic response to microenvironmental cGAMP (Figure 3D).

**Figure 3:**
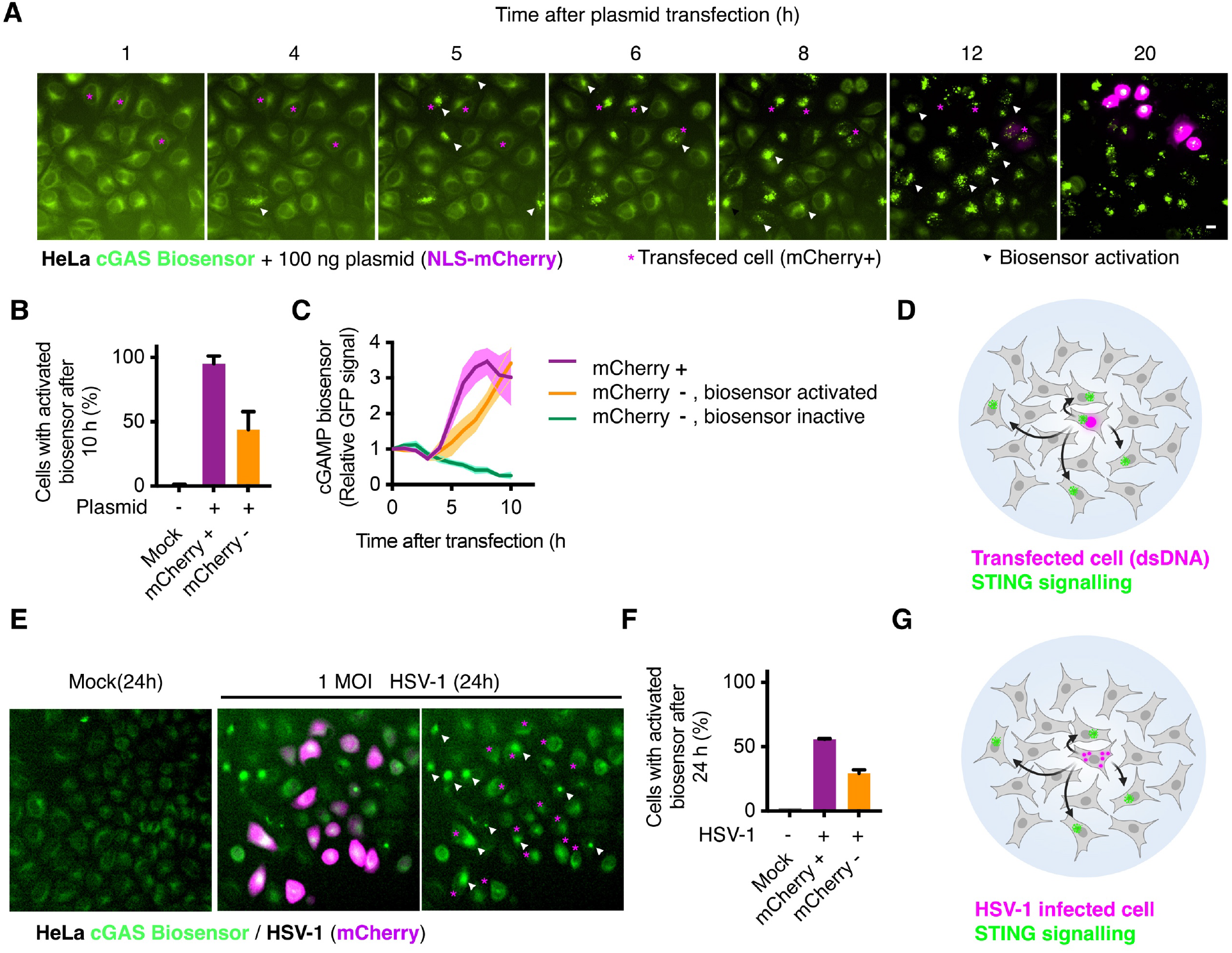
Monitoring foreign dsDNA with a novel cGAMP biosensor. **A-C**, Live cell imaging analyses of HeLa cGAMP biosensor cells transfected with a circular plasmid expressing SV40-mCherry (nuclear localization). In (**A**), magenta asterisks mark cells that express mCherry during the 24-hour recording, and white arrows mark the time-point in which cells first showed activation of their cGAMP biosensor. In (**C**) cells from the transfected condition are showed in 3 groups: Cells that will become mCherry positive and therefore were transfected with the plasmid, cells that did not express mCherry during 24 hours and showed activation of the biosensor during the first 6 hours, and cells that neither expressed mCherry during 24 hours nor showed activation of the biosensor during the first 6 hours. **D**, Suggested model of the spread of microenvironmental cGAMP-driven STING activation upon cytoplasmic dsDNA **E**,**F**, Live cell imaging analyses of HeLa cGAMP biosensor cells mock treated or infected with 1 MOI HSV-1 expressing mCherry, and monitored in an IncuCyte. In (**E**), magenta asterisks mark infected cells (mCherry positive) during the 24-hour recording, and white arrows mark cells first with clearly activated cGAMP biosensor. **G**, Suggested model of the spread of microenvironmental cGAMP and STING activation in a cell population upon HSV-1 infection.

To test the response of the biosensor to DNA viruses, we exposed HeLa cells to HSV-1 particles expressing mCherry (MOI = 1). Infected HeLa cells (mCherry positive) display a heterogenous, and largely low, STING signalling response as showed by the cGAMP biosensor (Figure 3E,F). Intriguingly, the biosensors at uninfected neighbouring cells display a quick and high response to the population infection. These results support previous findings showing that viral infections can hamper the intrinsic cellular STING response, thereby leading to innate immune evasion ^39,40^, and highlights that microenvironmental spread of cGAMP and/or released dsDNA to the neighbouring cells can be a source of STING activation in an infected cell population (Figure 3G).

Given the reliability of the biosensor to be activated by agonists (Figure 2A,B), these results highlight the suitability of this novel tool to monitor the heterogenous innate immune response via cGAS-STING signalling towards foreign DNA, especially in the context of viral infections.

### Monitoring host cytoplasmic dsDNA with a novel cGAMP biosensor

To determine the response of the biosensor to intrinsic sources of dsDNA, we performed apoptosis and chromosome missegregation experiments.

We induced apoptosis in HeLa cells using a cocktail of BH3-mimetics (see methods) that induce BAX/BAK activation by specifically blocking their inhibitors and monitored the kinetics of cell death for 24 hours in an IncuCyte plate reader using the internalization of DRAQ7 as a proxy. BAX/BAK activation induces the opening of the apoptotic pore and the permeabilization of the mitochondrial outer membrane, which is followed by release of mtDNA into the cytosol ^20,21,41^. In agreement with previous studies showing that activated apoptotic caspases cleave and inactivate cGAS thus preventing the engagement of the innate immune response in response to cytosolic mtDNA ^20,22,23^, triggering of apoptosis in HeLa cells resulted in cell death, but not activation of the cGAMP biosensor (Figure 4A-C). Importantly, inhibition of caspases activity with the pan-caspase inhibitor QVD-OPH (QvD) not only prevented cell death (Figure 4C), but also led to the activation of the cGAS biosensor, which was detectable 5 hours after treatment (Figure 4B). To confirm that the biosensor indeed responded to released mtDNA during apoptosis and not to other activators, we treated HeLa cells with 2’,3’-dideoxycytidine (DDC) for 1 week, which removes mtDNA by preventing its replication (Figure 4D) ^42^. In contrast to untreated cells, DDC-treated cells did not activate the cGAMP biosensor during apoptosis, including in the presence of the caspase inhibitors (Figure 4E,F). These results indicate that the efflux of mtDNA leads to cGAS-dependent activation of the cGAMP biosensor and support its reliability to explore e.g., the dynamics and signalling consequences of mitochondria outer membrane pores during apoptosis.

**Figure 4:**
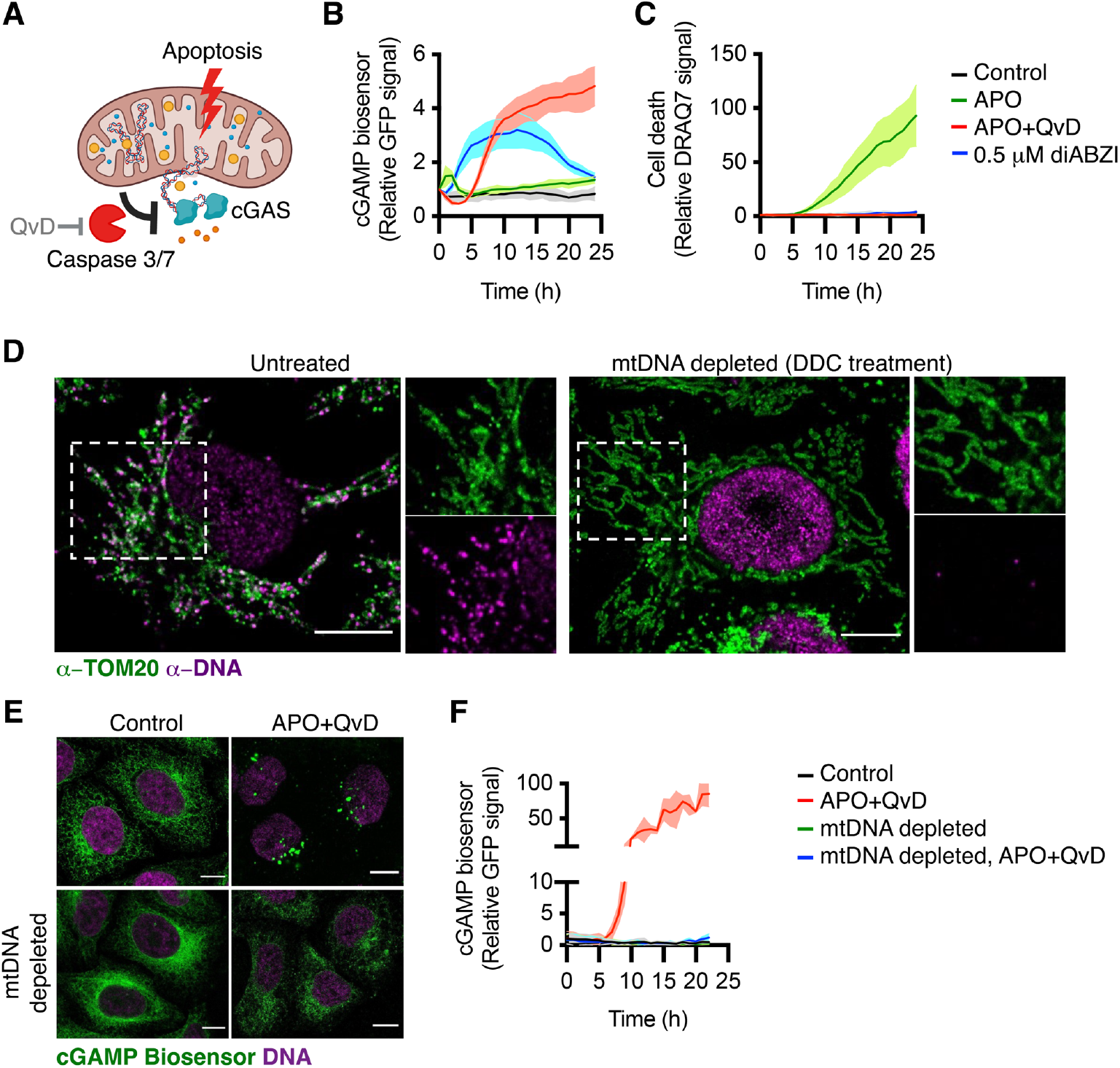
Monitoring mtDNA flux upon apoptosis with a novel cGAMP biosensor. **A**, Model of the caspase-dependent inhibition of cGAS during apoptosis. **B**,**C**, Cell population imaging analyses of HeLa cGAMP biosensor cells in IncuCyte in the presence or absence of an apoptotic cocktail of BH-3 mimetics, diABZI, with or without the caspase inhibitor QvD. In **(C**), cell death was quantified by cell permeabilization towards DRAQ5. **D**,**E**, Immunofluorescence experiments in HeLa cGAMP biosensor cells after 2-week treatment with DDC, or mock treated (Control). **F**, IncuCyte cell population imaging analyses of HeLa cGAMP biosensor cells iterated with DDC for 2 weeks (mtDNA depleted) or mock treated (Control) in the presence or absence of an apoptotic cocktail and the caspase inhibitor QvD.

Chromosome segregation errors such as lagging chromosomes can lead to formation of micronuclei ^43^. Rupture of the micronuclei’s nuclear envelope has shown to trigger cGAS recruitment ^24,26^, and it has been proposed as a critical mechanism of surveillance in the context of chromosomally instable (CIN) tumours ^44^. To examine this question, we generated mCherry-H2B and cGAMP biosensor HeLa cells and treated them with the MPS1 inhibitor reversine ^45^, which blocks the spindle assembly checkpoint thereby resulting in whole-chromosome missegregation and micronuclei.

We treated HeLa cGAMP-biosensor H2B-mCherry cells with 0.5 μM reversine, and monitored i) chromosome missegregation and micronuclei formation during the first 3 hours, and ii) biosensor activity during 24 hours by live cell imaging. Unexpectedly, HeLa cells harbouring micronuclei after mitosis did not show activation of the biosensor during the following 17-24 hours of the recording (Figure 5A,B). Given that a large proportion of micronuclei might not rupture during the recording, we performed immunofluorescence analyses in HeLa cGAMP biosensor cells after 18 hours treatment with reversine, which resulted in ∼33% of cells with cGAS^-^ micronuclei (likely intact) and ∼6% with cGAS^+^ micronuclei (ruptured) (Figure 5C-E). Consistent with their whole-chromosome segregation origin, 92% of micronuclei were positive for the centromere protein CENPC (Figure S2A,B). Intriguingly, HeLa cells with cGAS^+^ micronuclei did not display activation of the cGAMP biosensor compared to control cells treated with diABZI (Figure 5C,F), and other sources of cytoplasmic dsDNA (Figures 3-4).

**Figure 5:**
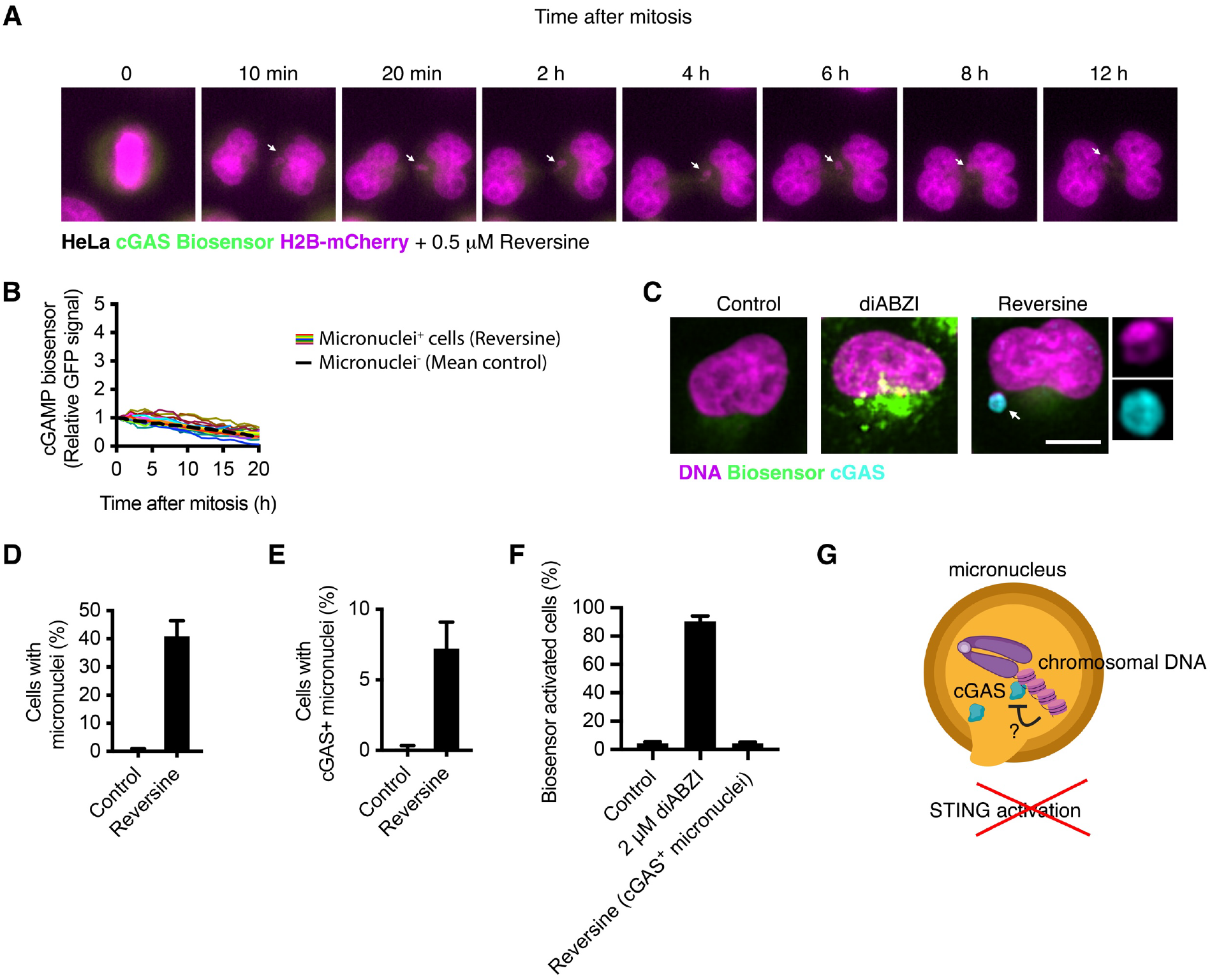
Micronuclei do not activate the cGAMP biosensor. **A**,**B**, Live cell imaging analyses of HeLa cGAMP-biosensor H2B-mCherry cells treated with 0.5 μM reversine, undergoing chromosome missegregation and micronuclei formation during the first 3 hours of the recording, and monitored for 17-20 hours. In (**A**), white arrows mark a lagging chromosome and the resulting micronucleus. **C-F**, Immunofluorescence experiments in HeLa cGAMP biosensor cells treated with 0.5 μM reversine for 18 hours. In (**F**) activation of the biosensor was scored as in previous analyses showed in Figure 2B and 3B. **G**, Ruptured micronuclei do not activate the cGAMP biosensor.

Taken together, these results indicate that the cGAMP biosensor is suitable to monitor host cytopismic dsDNA such as mtDNA flux during apoptosis, and suggest that ruptured micronuclei do not *per se* activate STING signalling, possibly due to cGAS inhibition by TREX1 and/or sequestration by the nucleosome histones H2A/B ^46 47,48^.

## DISCUSSION

Here, we presented a fluorescent cGAMP biosensor that can robustly react within 45 minutes and in a time- and concentration-dependent manner to STING agonists and cGAMP. We demonstrate that the cGAMP biosensor is suitable for single cell characterisation of the innate immune signalling dynamics upon foreign DNA, including viral infection spreading in a cell population. We suggest that this biosensor could be suitable to study mechanisms of innate immune evasion by viruses ^39,40^, as well as their recognition, including signalling spread through a complex cell population. Among other sources of cytoplasmic dsDNA, we also show that the cGAMP biosensor serves to monitor mtDNA release and signalling blockage by caspases following apoptosis. Given its robustness, this biosensor could help in further studying the contribution of mtDNA to inflammatory signalling associated with disease under different conditions, like mitochondrial dysfunction, cancer, and neurodegeneration ^49^.

Unlike mtDNA released to the cytoplasm, we show that cGAS recruited to ruptured micronuclei failed to activate the cGAMP biosensor. Our results are consistent with recent reports showing that both TREX1 and the nucleosome histones H2A/B inhibit cGAS activation towards genomic DNA ^46 47,48^, and suggest that other cytosolic pattern recognition receptors may underlie the interferon response upon chromosomal instability.

Finally, we showed that the biosensor sensitivity is sufficient to report microenvironmental cGAMP, including upon viral infection. We found that unlike permeable STING agonists or cytoplasmic dsDNA, microenvironmental cGAMP induce a heterogenous STING signalling response, suggesting the different cell intrinsic or extrinsic factors modulate the import, transport and/or stability of this immune-transmitter, with relevance for future studies on how the innate immune response spreads through tissues.

Taken together, we provide a tool to i) generate reporter cell lines in a single step; ii) easily monitor the activation cGAS-STING signalling pathway by live cell imaging, plate reader, or immunofluorescence analyses; and iii) capture the spatio-temporal and heterogenous dynamics of the response to cGAMP at single cell resolution.

## METHODS

The detailed materials and methods can be found at the end of the merge manuscript.

## ETHICS DECLARATIONS

The authors declare no competing interests.

## ACKNOWLEDGMENTS

We thank M. Knop for sharing reagents and cells. We thank the Nikon Imaging Center and the FACS facility at the University of Heidelberg for access to microscopes, cytometers, and for technical help. This work was funded by the Deutsche Forschungsgemeinschaft (DFG, German Research Foundation) and Heidelberg University through the Excellence Initiative – Explorer programme. S.S. holds a PhD fellowship from Boehringer Ingelheim Fonds.

## AUTHOR CONTRIBUTIONS

S.S, S.D.M-V, S.P.A. conceived, performed and analysed experiments. S.D., R.M., and D.I. performed and analysed experiments. G.P. contributed new reagents/analytic tools and supervised aspects of the project. S.P.A. designed the research, supervised the project and wrote the paper with input of the other authors.

## Supplementary

**Figure S1:**
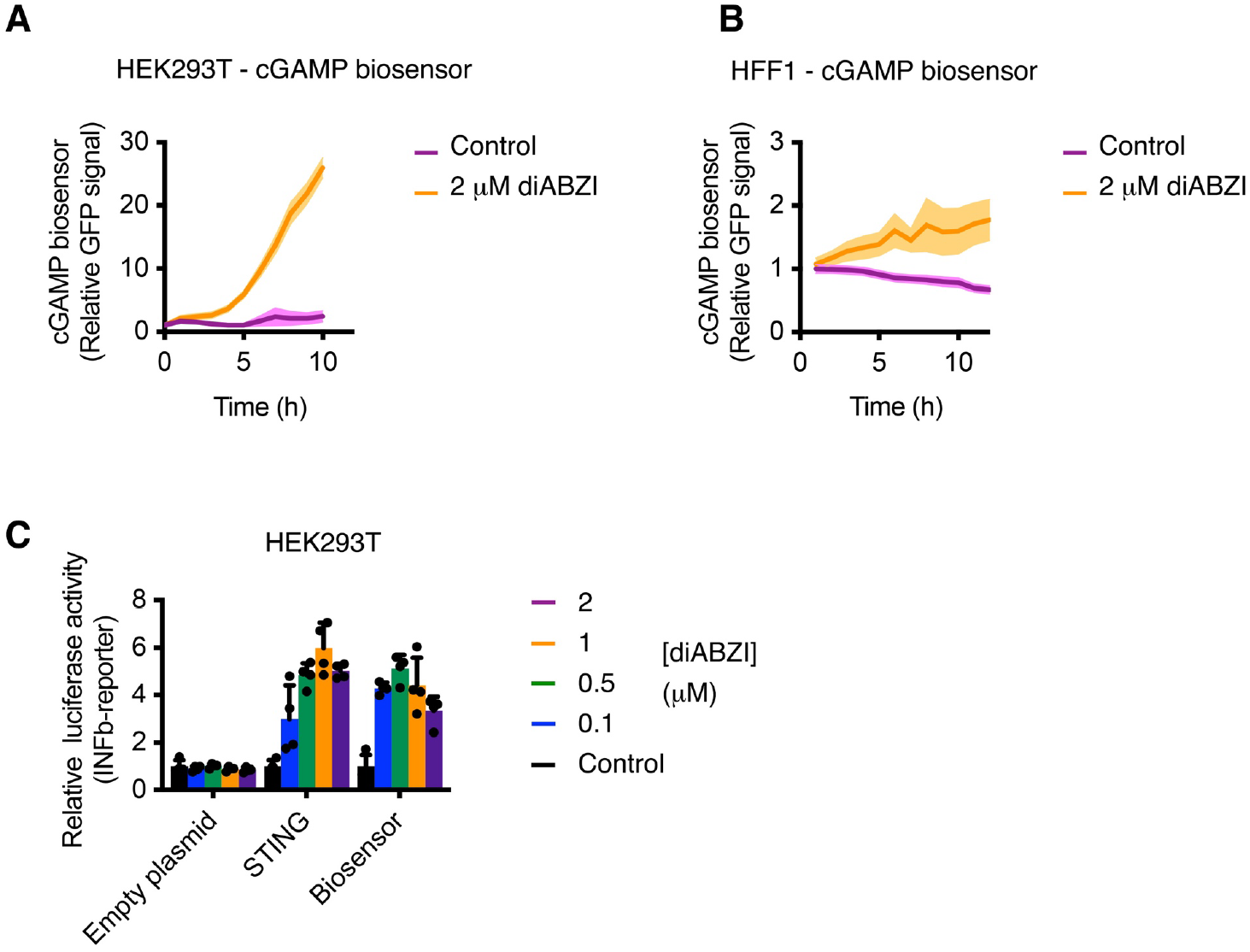
Validation of the cGAMP biosensor in HEK293T and HFF-1 cells. **A**,**B**, Live cell imaging analyses of HEK293T and HFF-1 cGAMP-biosensor cells treated with 2 μM diABZI. **C**, IFNb-Luc reporter assays in HEK293T cells transfected with empty plasmid or wt STING, or HEK293T cGAMP-biosensor cells, and treated with the indicated concentrations of the STING agonist diABZI. Note that HEK293T do not express endogenous cGAS or STING.

**Figure S2:**
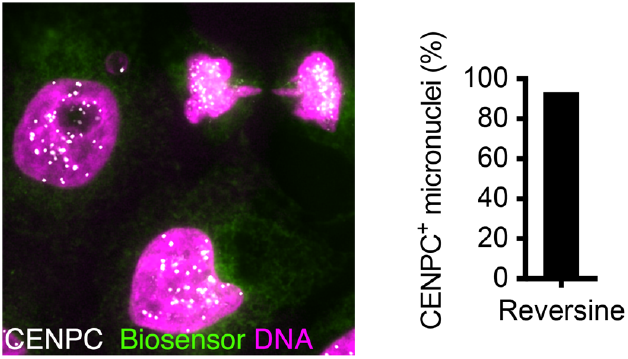
Characterisation of Micronuclei upon reversine in HeLa cells. **A**,**B**, Immunofluorescence experiments in HeLa cGAMP biosensor cells treated with 0.5 μM reversine for 18 hours and stained as indicated.

## SUPPLEMENTARY METHODS

### Constructs

The cGAMP biosensor utilises tandem split GFP ^50^. In detail, a construct containing STING-GFP11_x3_ (GFP(N)) and IRF3-GFP1-10 (GFP(C)) were cloned into a PiggyBac separated by a ribosome skipping peptide (P2A) to facilitate stochiometric expression (Referred from here on as cGAMP biosensor).

### Cell culture

HeLa and HEK293T cells (ATCC) were cultured in DMEM medium (Gibco) supplemented with 10% FBS and 1% penicillin/streptomycin.

To generate the HeLa and HEK293T cGAMP biosensor cells, the cGAMP biosensor was cloned into the PiggyBAC backbone plasmid (Biocat PB510B-1-SBI). The cGAMP biosensor plasmid was cotransfected with the PiggyBAC transposase plasmid (Biocat PB210PA-1-SBI) at a ratio of 500 ng to 100 ng respectively in a six well plate. Transfection was carried out with Lipofectamine3000 (ThermoFisher) according to manufacturer’s protocol. Cells were selected with 2 μg/ml and 3 μg/ml puromycin for the HeLa and HEK293T cell lines respectively. Monoclonal cell lines were then expanded from the surviving cells.

To generate the HeLa H2B-mCherry cGAMP biosensor cells, H2B-mCherry was cloned into the PiggyBAC backbone plasmid (Biocat PB510B-1-SBI). The H2B-mCherry plasmid was then co transfected with the PiggyBAC transposase plasmid (Biocat PB210PA-1-SBI) at a ratio of 500 ng to 100 ng respectively in a six well plate. Transfection was carried out with Lipofectamine3000 (ThermoFisher) according to manufacturer’s protocol. After 72 hours, the H2B-mCherry positive cells were pooled together via cell sorting and then monoclonal cell lines were generated from the polyclonal cells.

Where indicated, mtDNA depletion was achieved via 2’,3’ dideoxycytidine (DDC) treatment ^42,51^. Cells were growing in complete media with 1 mM sodium pyruvate and 40 μM of DDC for 6 days. Media was replaced every second day and passaged if needed. After treatment cells were collected for experiment.

Constructs and cell lines are available at request.

### Live cell imaging

HeLa and HEK293T cells stably expressing the cGAMP biosensor, H2B-mCherry, and/or HNRP-mCherry were seeded in a μ-slide 8-well chamber precoated with ibiTreat (Ibidi). Xy positions were first predetermined and then treatments were added before live cell imaging experiment was initiated. Live cell imaging was performed using an automated Nikon Eclipse Ti2 inverted microscope equipped with a 20x dry objective (NA 0.95) and a Nikon DS-Qi2 high-sensitive CMOS monochrome camera. Multipoint acquisition was controlled by NIS-Elements 5.1 software. Image stacks were recorded every 1 hour or 15 minutes for up to 18-24 hours or 2 hours respectively in an OkoLab environmental chamber at 37 °C and 5% CO2. Images were analysed using ImageJ 2.0.0 software. In particular, biosensor fluorescence signal was tracked manually and monitored as median fluorescence insensitive (MFI). Relative signal was calculated by subtracting the background signal and dividing the intensity of all time points by the first frame.

For the apoptosis analyses in Figure 3, HeLa cGAMP biosensor cells were performed using IncuCyte S3 (Sartorius) at 37°C 5% CO 2. For this, cells were seeded in a 96-well plate. Next day, cells were treated with either vehicle or BH-3 mimetic drugs ABT-737 (Hölzel;10 μM) and S63845 (10 μM) to induce MOMP/apoptosis in presence of pan-caspase inhibitor (QvD; 10 μM). STING agonist, diABZi was used as positive control for STING activation. After adding treatments, plate was inserted in IncuCyte chamber and 3 images per well were acquired every 1 h for 22 h. Images were analyzed using IncuCyte analysis software module. Green positive object per mm^2^, information was used to create the graph.

For the infection analyses in Figures 4, HeLa cGAMP biosensor cells were seeded at a density of five thousand cells per well in a 96-well plate. Next day, cells were infected with HSV-1 mCherry (Multiplicity of infection, MOI = 1), in 2% FCS DMEM. STING agonist, diABZI, was used as positive control for STING activation. After treatment, cells were placed in an IncuCyte SX5 (Sartorius) at 37°C, 5% CO_2_ system, and imaged every 2 h for 3 days. Images were analyzed using IncuCyte analysis software module.

### DNA and siRNA transfection

Cells were transfected with 25nM or 50 nM siRNAs (Sigma) using Lipofectamine RNAiMAX transfection reagent (Thermo Fisher Scientific 13778075). 48-72 h after transfection, cells were further analyzed. The following siRNAs were used: siRNA against Scramble (SIC001) or an siRNA pool against firefly luciferase mRNA (U47296) (Dharmacon D-001206-14-20) as negative controls, and an siRNA pool against TBK1 (Dharmacon M-003788-02-0010).

For the circular plasmid transfection analyses, cells were transfected with 100 ng of PCS2+ containing SV40-mCherry per well of a a μ-slide 8-well chamber precoated with ibiTreat (Ibidi) using X-tremeGENE 9 DNA transfection reagent (Roche) immediately before imaging.

### Immunofluorescence

HeLa cells were seeded on coverslips in 12-well plates. Where indicated, cells were first transfected with siRNA for 48-72 h. Cells were fixed with 2% paraformaldehyde in PBS and stained, according to the experiment, for DAPI and the indicated proteins using guinea pig anti-CENP-C (MBL PD03) and rabbit anti-cGAS (CST 79979) and then probed with donkey anti-guinea pig Cy3, donkey anti-rabbit Alexa594 (ThermoFisher A21206) or donkey anti-rabbit Alexa647 (Thermofisher A31573).

Coverslips were imaged with a Nikon Eclipse Ti using a Nikon Plan Apo λ 60x NA 1.40 with oil immersion and the NIS Elements software. Data was analyzed using ImageJ 2.0.0.

For the experiments in Figure 4, cells were seeded on glass coverslips, treated with diABZI, vehicle or BH-3 mimetic drugs to induce MOMP/apoptosis, fixed in 4% paraformaldehyde and blocked with BSA. Cells were immunostained for anti-DNA (1:200, Progen-690014S) and anti-TOMM20 (1:200, Atlas-HPA011562) and secondary antibodies used were Abberior STAR ORANGE and STAR-RED (Abberior, 1:200 diluted). Images were taken in the Abberior confocal microscope and processed in ImageJ.

### Western blotting

For Western blotting, cells were lysed in full lysis buffer (50 mM Tris-HCl, pH 7.5, 150 mM NaCl, 1% Triton X-100, 0.05% SDS, 1 mM β-mercaptoethanol, 2 mM EDTA, 1x protease phosphatase inhibitor cocktail (Thermo Fisher)). The cleared lysates were mixed with 4x NuPAGE loading buffer, resolved on 10% NuPAGE gels and transferred to nitrocellulose membranes. For western blot experiments the following antibodies were used: mouse anti-α-tubulin (Sigma Aldrich, T9026), rabbit anti-TBK1 (CST 38006), rabbit anti-STING (CST 13647), rabbit anti-phospho-TBK1 (CST 5483), and rabbit anti-phospho-STING (CST 50907). Membranes were then probed with goat anti-mouse IgG HRP (Millipore AP308P) or goat anti-rabbit IgG HRP (CST 7074).

### Image data processing

Raw images were imported to Fiji (ImageJ, v2.0) prior to their export to Photoshop 2020 for panel arrangement. Linear changes in contrast or brightness were equally applied to all controls and across the entire images. The models and schemes were created with BioRender.com.

### Statistical analyses

Data are shown as mean with standard error of the mean (SEM or SD), as indicated in the figure legends. Where indicated, Student’s t-tests (two groups) or one-way ANOVA analyses with Tukey correction (three or more groups) were calculated using Prism v8. Significance is indicated as: *P < 0.05, **P < 0.01, ***P < 0.001, or n.s.: not significant.

